# Birdsong Phrase Verification and Classification Using Siamese Neural Networks

**DOI:** 10.1101/2021.03.16.435625

**Authors:** Santiago Rentería, Edgar E. Vallejo, Charles E. Taylor

## Abstract

The process of learning good features to discriminate among numerous and different bird phrases is computationally expensive. Moreover, it might be impossible to achieve acceptable performance in cases where training data is scarce and classes are unbalanced. To address this issue, we propose a few-shot learning task in which an algorithm must make predictions given only a few instances of each class. We compared the performance of different Siamese Neural Networks at metric learning over the set of Cassini’s Vireo syllables. Then, the network features were reused for the few-shot classification task. With this approach we overcame the limitations of data scarcity and class imbalance while achieving state-of-the-art performance.

## 1 Introduction

Current evolutionary studies of bird vocalizations require *automatic unit segmentation and classification* methods capable of generalizing not only across species but also dealing with noisy environments. By extracting patterns from large-scale recordings new hypotheses regarding birdsong structure could be tested while at the same time reduce human bias and increase research reproducibility. Previous work in this area has been done with different species using a wide range of techniques such as Hidden Markov Models (Kaewtip, Taylor, & Alwan, 2016; Koumura & Okanoya, 2016), Support Vector Machines (Arriaga, Kossan, Cody, Vallejo, & Taylor, 2013), Dynamic Time Warping (Tan, Alwan, Kossan, Cody, & Taylor, 2015) and Deep Learning (Koops, van Balen, & Wiering, 2015). Within these lines in this work we focus on *automatic unit classification* by comparing different Siamese Neural Networks, a few-shot machine learning technique capable of discriminating syllable classes from scarce data.

Our main objective is to find a classifier capable of dealing with data sparsity and class imbalance in *Vireo cassinii* syllable repertoires. These are common issues in birdsong research given limited recordings and the existence of rare syllables. In the long run, we expect our results to have a significant impact in ecology and evolutionary biology, where new analysis tools will overcome the limitations of manual (human) recognition (Kershenbaum et al., 2016). We propose a few-shot machine learning approach to classify *Vireo cassinii* syllables using a family of siamese neural networks with different encoders, including convolutional, LSTM, Bidirectional LSTM fully-connected and “encoderless” siamese networks. The latter is equivalent to *k*-nearest neighbor classifier with euclidean metric.

The main contribution to biology is providing a tool to increase our knowledge about sophisticated signaling strategies and syntactic structures in non-human species as birds. By the other side, the model is computationally interesting since few-shot classification approaches constrain the algorithm to learn to discriminate among instances by only observing a few samples from each class. This is similar to the kind of learning observed in children, which develop sophisticated rules about new word categories from very few or even no examples at all (Yip & Sussman, 1997; Furbee, 1992). In contrast, most Deep Learning models rely on large data sets to achieve acceptable performance. As far as we know, few-shot deep learning has not been applied to birdsong, although in principle the general approach can be replicated for almost any modality or domain.

### 1.1 Birdsong structure

Acoustic sequences are ubiquitous, from bird songs to human speech and music. More often than not they convey meaning, and have an important role in evolution as individuals can take advantage of the information contained in them (Kershenbaum et al., 2016). But when a bird sings how do we know whether communication has occurred? It is generally held by biologists that if the signal modifies the behavior of the receiving animal, then we can infer that communication has taken place. Similarly, we might say an acoustic sequence carries information when it has the potential to reduce uncertainty on the part of the receiver.

Bird acoustic sequences, also known as vocalizations, can be divided into *songs* and *calls*. In general songs tend to be long, complex vocalizations mostly produced by males during the breeding season for maintenance of territories and mate attraction. To these features there are innumerable exceptions covered by (Kershenbaum et al., 2016). By the other side, calls tend to be shorter, simpler in structure and produced by both sexes throughout the year. They are less spontaneous than songs and are usually related to specific functions such as flight, threat and alarm. One of the great questions of ornithology is why *Passerines* have evolved such complex songs and a special neural pathway to learn them. A question that can be tackled through algorithmic segmentation and classification methods (Catchpole & Slater, 2008).

*Songs* are subdivided in *phrases, syllables* and *elements*. Each *phrase* consists of a series of units (*syllables*) which occur together in a particular pattern. Similarly, *syllables* when complex, can be constructed from several of the smallest building blocks of all, known as *elements*. Regarding *songs*, each bird can have more than one version, making up a *repertoire* of song types. It is important to mention in the literature *phrases* and *syllables* are used interchangeably but ultimately they refer to medium-sized fragments of bird vocalizations.

### 1.2 Birdsong analysis techniques

Accurate analysis of bird vocalizations depends on appropriately characterizing their constituent units. Nevertheless, there is no single definition of unit, these vary widely across researchers. This is by no means whimsical as the characterization of units depends on the question being addressed.

Often the details of acoustic production and perception are hidden from the researcher, in consequence definition of acoustic units has to be carried out on the basis of observed acoustic properties. The first step is defining what possible functions a sound has, then formulating appropriate hypotheses after a period of observation and field study, which relates the singing bird to its habitat, general life and evolutionary history. Having observed and listened to birds in their habitat, the next step is to make audio recordings of their song and analyze them (Catchpole & Slater, 2008). Most analytic methods for unit classification assume they can be divided into discrete, distinct categories (Clark, Marler, & Beeman, 1987). According to this hypothesis, Kershenbaum et al. (2016) describe four main approaches to classify units by their acoustic properties:

1. Manual classification Units are “hand-scored” by humans searching for consistent patterns in spectrograms or by listening sound recordings without the aid of a spectrogram. Even if humans are good at pattern recognition, manual segmentation and classification is time consuming and prevents taking full advantage of large acoustic data sets generated by automated recorders. Similarly, this difficulty hinders research reproducibility as sample sizes studied tend to be too small to draw firm conclusions (Kershenbaum, 2014). Furthermore, manual classification can be prone to subjective error, and inter-observer reliability should be used (and reported) as a measure of the robustness of the manual assessment (Kershenbaum et al., 2016).
2. Classification of manually extracted metrics An alternative to manual segmentation is using feature extraction, for example: duration, pulse repetition rate, spectral centroid, Mel Frequency Cepstral Coefficient (MFCC) among others. These features are then used in classification algorithms and mathematical techniques such as principal component analysis (PCA), discriminant function analysis or classification and regression trees (CART). In this category we can find semi-automatic techniques where features are extracted by standard algorithms and then verified by human analysts (Kershenbaum et al., 2016).
3. Fully automatic metric extraction and segmentation Automatic segmentation or recognition of acoustic units is not prone to inter-observer variability of manual classification. However, current implementations are: not generalizable to all species (Pearre, Perkins, Markowitz, & Gardner, 2017), very sensitive to hyperparameters (Ranjard & Ross, 2008), require pre-determined syllable classes and boundaries (Koumura & Okanoya, 2016) or struggle at recognizing subtle features that can be detected both by humans and birds (i.e. high false positives rate or low accuracy in test sets) (Fukuzawa, Marsland, Pawley, & Gilman, 2017; Koops et al., 2015). Interestingly, manual classification has been shown to out-perform automated systems in cases where the meaning of acoustic signals is known a priori, possibly because the acoustic features used by fully automated systems may not reflect the cues used by the focal species (Kershenbaum et al., 2016). Although, there is motivation in developing fully automated segmentation and classification algorithms given they allow large scale analysis of birdsong recordings.

The definition of a unit for a particular species depends on the question being addressed and is dependent on a large number of factors. In particular, availability of behavioral information, such as responses of individuals to playback experiments and morphological information. Kershenbaum et al. (2016) suggest the following protocol to define acoustic units:

1. Determine what is known about the production mechanism of the signalling individual.
2. Determine what is known about the perception abilities of the receiving individual. Perceptual limitations may substantially alter the structure of production units.
3. Choose a classification method, such as manual, semi-automatic, or fully automatic. Some unit types lend themselves more readily to certain classification techniques than others.

Various algorithmic approaches to birdsong recognition have been made in the last years. For example, Potamitis used Deep learning for detecting bird vocalisations (Potamitis, Ntalampiras, Jahn, & Riede, 2014). Ranjard and Ross achieved unsupervised bird song syllable classification using evolving neural networks (Ranjard & Ross, 2008). Modern fully automatic techniques rely on Hidden Markov Models and Convolutional Neural Networks trained on manually annotated data (Koumura & Okanoya, 2016).

### 2 Materials & Methods

We propose a few-shot machine learning approach to classify *Vireo cassinii* syllables using a family of siamese neural networks with different encoders, including convolutional, LSTM, Bidirectional LSTM fullyconnected and “encoderless” siamese networks. The latter is equivalent to *k*-nearest neighbor classifier with euclidean metric.

We studied the generalization capabilities of this family of siamese models under 1,3,5 and 7 examples. In order to do so, each model was trained to carry out a verification task, which then generalized to few-shot classification: First, it learned to assign a low distance score to pairs of syllables of *Vireo cassinii* belonging to the same class. Then, we used the learned similarity function to evaluate syllables in a pairwise manner against a representative instance of each class (i.e. average of training set). The pairing with the lowest distance score was awarded the highest probability for the classification task. Figure 1 depicts the Siamese Network model in general.

**Figure 1:**
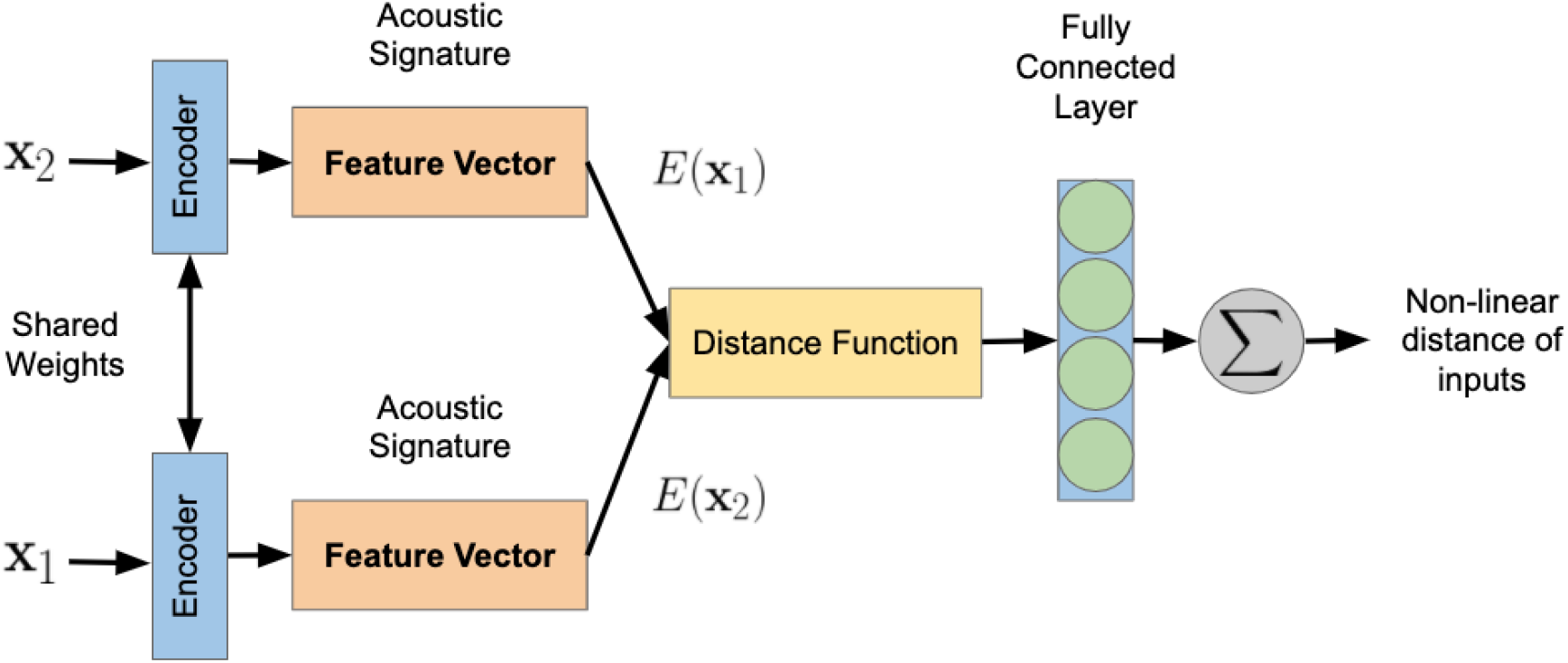
Siamese Network Model Overview

### 2.1 Species: *Vireo cassinii*

This species, also known as Cassin’s Vireo (abbreviated as “CAVI” in singular, or “CAVIs” in plural in this paper) belongs to the order of *Passeriformes* and to the *Vireonidae* family. It is commonly found in many coniferous and mixed-forest bird communities in far western North America. Only the males of this species give full songs, and their songs have been described as a jerky series of blurry phrases, separated by pauses of ≥ 1 second. Each phrase is made up of 2 to 4 notes (syllables), with song often alternating between ascending and descending phrases. The song is repeated tirelessly, particularly when the singing male is unpaired (Goguen & Curson, 2002).

Songs from two males on two different territories in a conifer-oak forest in California were recorded at approximately 800 m elevation (38°29′04′*N*), (120°38′04′W), near the city of Volcano in California (Amador County), USA. The data collection was done between April and June 2010. Manual inspection was done using Praat software to identify the phrase class, and mark the start and end times of each song element. The recordings and annotations for this 2010 collection are freely available online at Bird-DB. This was the same data set used by Tan, Kossan, Cody, Taylor, and Alwan (2013) for bird phrase verification and classification with a sparse representation-based classifier. In this work it will be referred as *Tan2013* data set.

### 2.2 Data & Tools

In *Tan2013* dataset there are 1116 tokens in total grouped in 64 classes, with a range of 1 to 73 tokens each (See Figure 7). The more frequently observed 32 phrase classes have at least *n* = 12 tokens. These conform the *filtered set*, which amounts to 1033 tokens. Phrases depicted in Figure 8, were used as a *support set* for siamese neural network classification. Since our main interest was to test siamese neural networks performance in a *k*-shot scenario with *k* = [1, 3, 5, 7], *n* − *k* tokens from each of the 32 classes were removed at random from the *filtered set* (See Figure 5) to make the *test set*. The remainder conforms the *training set*. Infrequent classes amount to 83 tokens, and are one of the main reasons few-shot learning approaches are relevant to birdsong research.

Regarding phrase duration, the longest instance is of class ‘at’ with 27525 audio samples, and the shortest is of class ‘bm’ with audio 3794 samples. With a sample rate of 22.5 Khz they are respectively 1.24 s and 0.172 s long. In the *filtered set* the shortest is the same but the longest is of class ‘ac’ with a duration of 1.06 s and 21352 audio samples (See Figure 6).

### 2.3 Database: Bird-DB

Projects on the acoustic monitoring of animals in natural habitats generally face the problem of managing extensive amounts of data produced for experimentation. While there are many publicly accessible databases for birdsong recordings, such as Xenocanto (Planqué & Vellinga, 2005) and Macaulay Library from Cornell University, most of them lack annotated song sequences.

Bird-DB provides an interface and annotated database for studying the syntax of bird song. Users are capable of selecting attributes relating to several general aspects of the stored recordings, for instance: recording hardware, location and environment. Queries can be narrowed down to specific species and individuals. The database returns a list of records meeting those criteria, with links to the appropriate audio and annotation files in *TextGrid* format (Arriaga, Cody, Vallejo, & Taylor, 2015).

### 2.4 Pre-processing

Phonological analysis of animal vocalizations requires specialized software providing visualization, annotation and measurement tools for analyzing audio recordings of any length. *Raven Pro* (Bioacoustics Research Program & Program, 2014), a software maintained by The Cornell Lab of Ornithology, is widely used in bioacoustic research. Nevertheless, since the recordings from Bird-DB were annotated in *Praat* software (Boersma & Weenink, 2011), we stick to this option. Both have similar specifications, despite *Praat* was designed for human phonetics analysis, whereas *Raven Pro* emerged from the broader field of bioacoustics.

Birdsong phrase annotations in Praat are processed and stored using the *Textgrid* format. Audio and annotation files are stored separately. A *TextGrid* object consists of a number of tiers of two kinds:

1. **Interval tier:** A connected sequence of labelled intervals, with boundaries in between.
2. **Point tier:** A point tier is a sequence of labelled points.

Through *Praat*’s interface users can store and label sequences of intervals, which later are employed to segment recordings. It is important to note birdsong phrase identifiers, like *bm* or *bp*, have no intrinsic meaning, they only provide a notation system to label classes of sounds.

Recordings were annotated by humans and segmented using a Python library (*praatIO*) by taking as an input the corresponding pairs of (*Textgrid*, audio file). The output of this procedure are a set of labeled audio (.wav) segments at a sample rate of 22,050 Hz and 32 bit resolution each. Since the meaningful information of *Tan2013* CAVI database is within the range of 1 kHz and 8 kHz, the rest of the frequencies can be safely removed using a bandpass filter. For this reason we applied a Butterworth Bandpass filter with 1 kHz and 8 kHz cut-off frequencies.

The sampling rate was first reduced to 20 kHz because energies of interest are below 10 kHz. Since every phrase instance has a variable duration, to generate a feature vector of the same dimension for each token, a file-duration-dependent frame shift was used to compute its spectrogram. The frame-shift was calculated by Tan (Tan, Kaewtip, Cody, Taylor, & Alwan, 2012) as follows for *t* = [0, *N* − 1], with *N* as the number of frames per token:

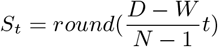

*D* and *W* denote respectively file duration and frame length in number of samples, whereas the starting sample index for frame *t* is denoted by *S*_*t*_. We used the parameters suggested by Tan (Tan et al., 2012), 64 frames per token and a frame length *W* of 20 ms, which amounts to 400 samples at a 20 kHz sampling frequency.

A 512-point Fast Fourier Transform is computed at each frame, and values are converted to decibels (dBFS) units. Given most of the bird phrase energy content falls within 1 and 8 kHz, only frequency bins corresponding to this range are retained. Finally, the sequence of spectrogram vectors is normalized per phrase.

### 2.5 Syllable Classification Models

We evaluated and implemented in Keras framework, four siamese networks with the following feature extractors: *convolutional (CNN), fully-connected (FCN), LSTM, bidirectional-LSTM* networks for *k* = [1, 3, 5, 7], these values were chosen for benchmarking purposes, as they were the ones reported by Tan et al. (2012). In addition to *Tan2013* nearest subspace approach, we compare the performance of siamese *k*-shot learners to a *zero neural network (Zero)*. This is a siamese neural network with no feature extractor mimicking the behavior of a *k*-nearest neighbor classifier with euclidean metric. As previously mentioned, the remaining *n* − *k* examples, where *n* is the number of items in each class, were used as a *test set*.

These five models were chosen for they are representative of the most used neural network architectures (Goodfellow et al., 2016). Furthermore, they embody different computational principles regarding temporal processing, parameter sharing, sparse interaction and connectivity. We summarize neural network details in Table 7 in the Appendix.

Given the stochastic nature of the sampling and optimization procedures in siamese neural networks, we ran each model 30 times and provide a 95% confidence interval for each value of *k*. In this way we can assess model sensitivity to training set size under normality assumptions. Hypothesis testing results are reported in the following chapter. The *support set* was generated by averaging per class the spectrograms in *training set*. Thus, the shape of input data is an array of shape (64, 128). The *support set* is used during classification as a set of exemplars to which distances are measured.

Even if it is desirable, we did not carry out cross-validation nor hyperparameter optimization due the high computational cost involved in training multiple deep neural networks. Experiments were performed on a laptop with a NVIDIA GeForce GTX 1050 GPU.

All pairs used to train siamese networks were sampled at random from the pre-processed *filtered set*, which amounts to 1033 instances across 32 classes. During each training round we sampled a total of 64000 pairs of instances, 2000 per class consisting of half same and half contrasting pairs. The test set contained 6400 pairs sampled at random, 200 per class. Finally, the output of siamese networks is interpreted as a metric between the input spectrograms of shape (64, 128). With this metric we carry out classification by assigning to each query the class of the closest instance in the *support set*.

## 3 Results

Does increasing the value of *k* (shots) improve the performance of few-shot models? To what extent computationally expensive models, such as Siamese Bidirectional LSTM benefit from having more training data (larger values of *k*)? To answer these questions we analyzed the average accuracy of different siamese models at four values of *k*. Average accuracy was computed over the most frequent 32 phrase classes. Significance across performance for each model was evaluated using pairwise z-tests for the difference between average classification test accuracy with different values of *k*.

Tables 1 to 5 provide data on the average classification accuracy of each model at the test set for different values of *k*. Since training sets were small (*k* = 1, 3, 5, 7), and models’ training phase relied on stochastic optimization techniques, we trained each model 30 times with different training set samples and random seeds. This decision was taken on the grounds of computational resources availability (i.e. experiments were performed on a mid-end personal computer) and the central limit theorem. For more details on the implementation please contact the author to obtain the scripts.

**Table 1:**
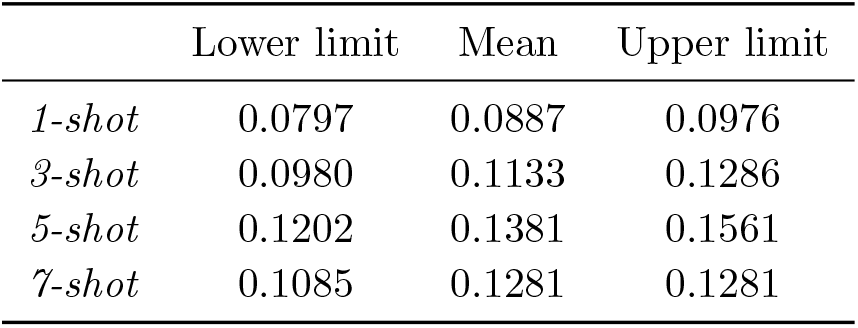
95 % confidence intervals for siamese FCN test classification accuracy

**Table 2:** 95 % confidence intervals for siamese CNN test classification accuracy

|  | Lower limit | Mean | Upper limit |
| --- | --- | --- | --- |
| *1-shot* | 0.5140 | 0.5396 | 0.5652 |
| *3-shot* | 0.7113 | 0.7324 | 0.7536 |
| *5-shot* | 0.7382 | 0.7570 | 0.7759 |
| *7-shot* | 0.7522 | 0.7703 | 0.7884 |

Fully-connected siamese network (FCN) did not benefit significantly from increasing values of *k*. FCN shows accuracy distributions with high variance, indicating high sensitivity to the selection of training instances per class. Highest classification accuracy obtained by this model was 12.81% with 7 shots. Convolutional siamese network greatly improved with respect to FCN, but p-values show diminishing returns after *k* = 3. Highest classification accuracy was 77.03% with 7 shots. LSTM siamese network had a similar performance (~ 73%) to its convolutional counterpart for *k* = 3, but overcame it at *k* = 5, 7, indicating greater learning capacity of LSTM siamese network. Highest accuracy for LSTM siamese network was 82.03% with 5 shots. Bidirectional LSTM siamese network beat all of the previous models with a mean accuracy of 85.14% using 3 shots. Highest accuracy was 91.31% with 7 shots. Finally, the Zero neural network beat all of the previous models excepting bidirectional LSTM with a mean accuracy of 83.17% using 3 shots. Highest accuracy of the Zero neural network was 90.10% with 7 shots, 1.21% below the bidirectional siamese network highest accuracy.

LSTM mean accuracy improves less significantly as *k* increases, as opposed to the bidirectional version. We can confirm a similar situation for the convolutional siamese network. Additionally, accuracy figures from the Appendix show higher overfitting for the convolutional model compared to LSTM and Bi-LSTM models for *k* = 7, as the gap between test and training conditions is larger. This might be explained by the fact LSTM models account for the sequential structure of spectrograms. We are faced with a low complexity model (*Zero Network*) performing as good as our most complex one (bidirectional LSTM). We will try to explain this in the Discussion section alluding to the *manifold hypothesis* and the structure of data after applying dimensionality reduction techniques.

## 4 Data Visualization

Figure 2 shows the behavior of the same data after applying Principal Component Analysis (PCA), a linear dimensionality reduction technique. We projected the *filtered set* on the first two principal components, those with the greatest accumulated variance.

**Figure 2:**
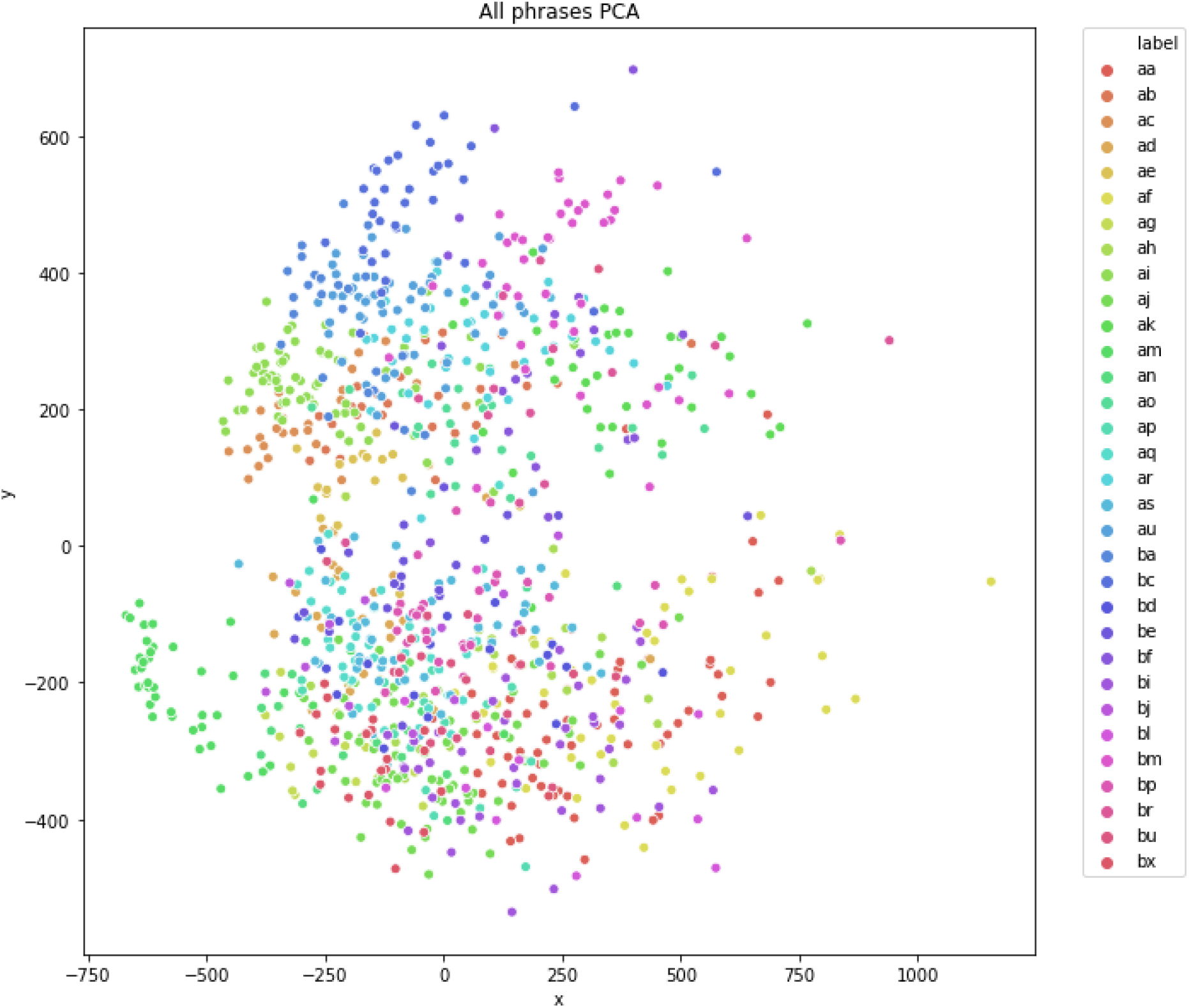
Phrases projected using the first two principal components

Figure 3 shows the behavior of the full dataset (i. e. *filtered set*) after applying t-Distributed Stochastic Neighbor Embedding (t-SNE), a dimensionality reduction technique minimizing the Kullback-Leibler divergence between the joint probabilities of the low-dimensional embedding and the high-dimensional data (Van Der Maaten & Hinton, 2008). The parameters we used were Perplexity: 25; Learning rate: 200; Metric: Euclidean; Dimension: 2.

**Figure 3:**
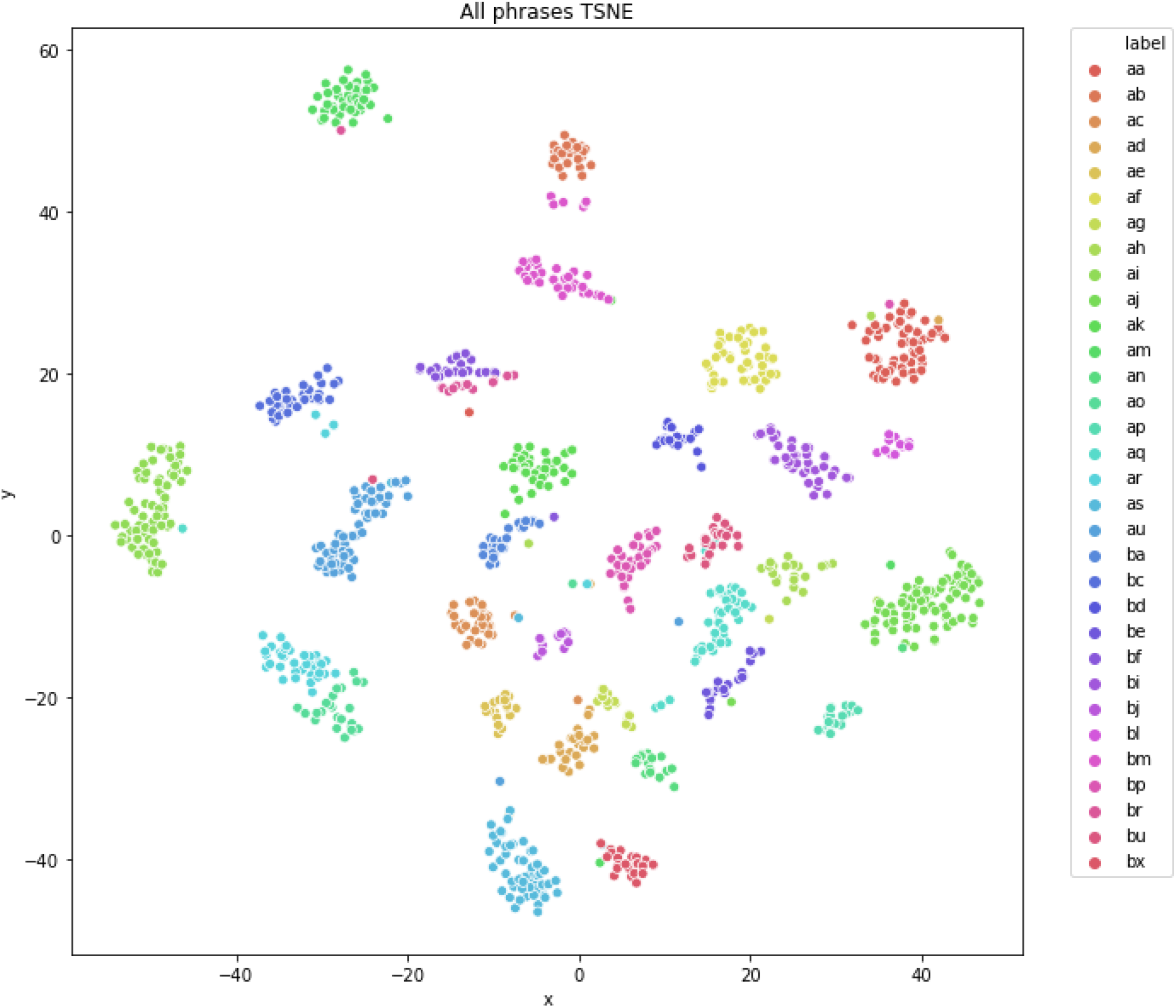
Phrases projected using t-SNE

Figure 4 shows t-SNE projected data along with the centroids computed from an arbitrarily selected support set with *k* = 7. Centroids might be interpreted as class prototypes around which most instances cluster. In all cases, before applying dimensionality reduction, spectrograms were flattened to obtain vectors of shape 1 × 8192. Then, these vectors were projected on a 2D plane.

**Figure 4:**
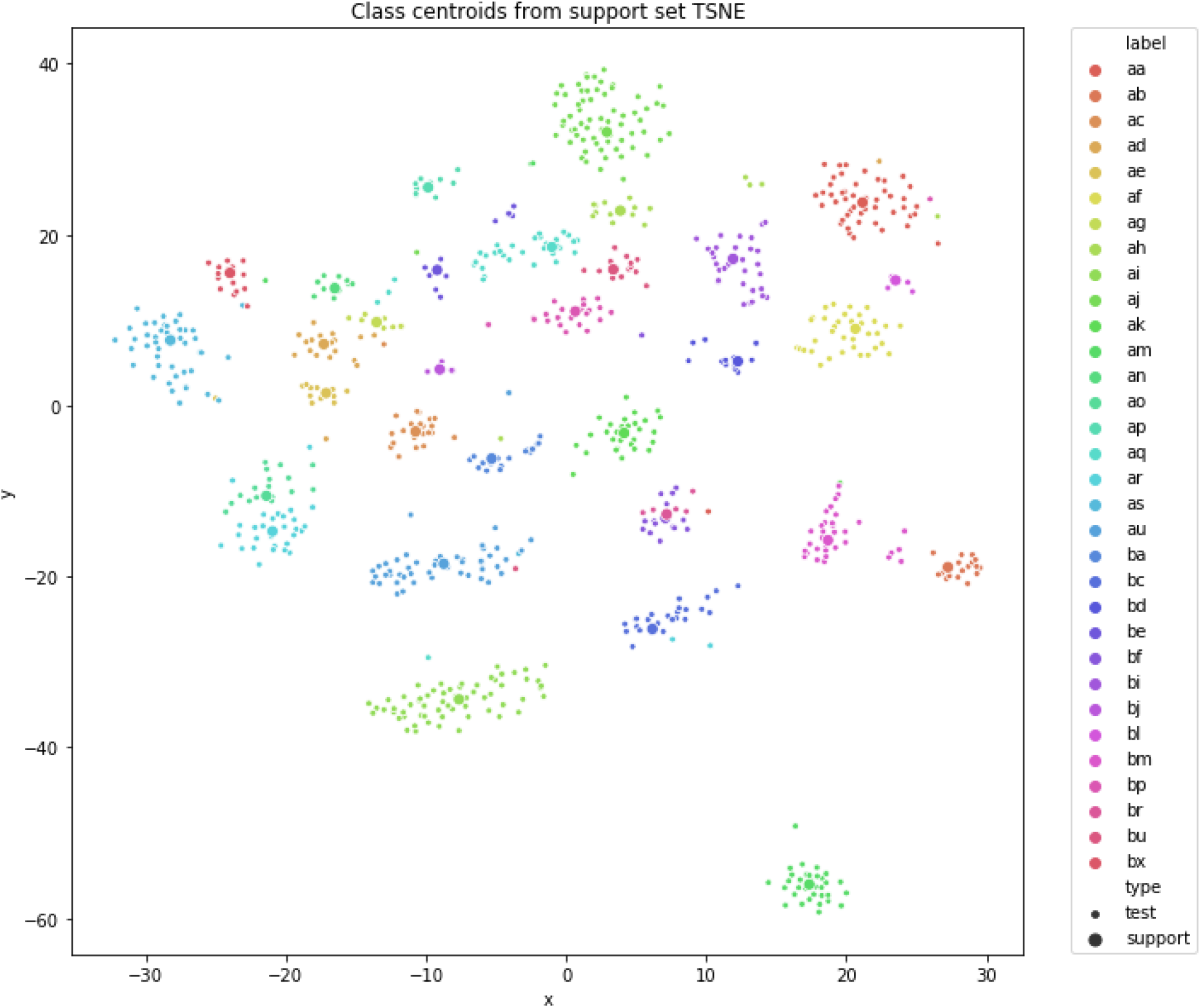
Test set and class centroids computed from support set

## 5 Discussion

The closest study to ours, by Tan et al. (2013), measured the performance of a sparse representation (SR) classifier for bird phrase verification and classification in the same dataset we used (referred as *Tan2013*). They found that when evaluated against nearest subspace (NS) and support vector machine (SVM) classifiers, the SR classifier had the highest test classification accuracy after dimensionality reduction with PCA: 89.6% using 7 shots. See Table 6.

In our study the Zero siamese network reached a similar test classification accuracy with 7 shots (90.10%), while the bidirectional LSTM reached it with 5 shots (90.24%). To date, for *Tan2013* dataset this has been the highest accuracy (see Tables 4 and 5). Therefore, our best models can be considered state of the art.

**Table 3:**
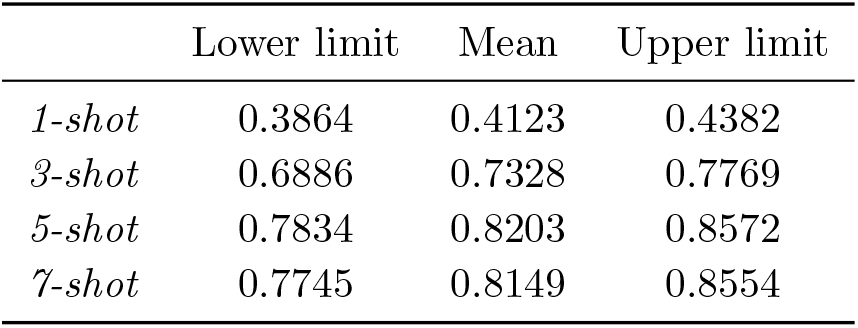
95 % confidence intervals for siamese LSTM test classification accuracy

|  | Lower limit | Mean | Upper limit |
| --- | --- | --- | --- |
| *1-shot* | 0.3864 | 0.4123 | 0.4382 |
| *3-shot* | 0.6886 | 0.7328 | 0.7769 |
| *5-shot* | 0.7834 | 0.8203 | 0.8572 |
| *7-shot* | 0.7745 | 0.8149 | 0.8554 |

**Table 4:** 95 % confidence intervals for siamese bidirectional LSTM test classification accuracy

|  | Lower limit | Mean | Upper limit |
| --- | --- | --- | --- |
| *1-shot* | 0.4710 | 0.4996 | 0.5282 |
| *3-shot* | 0.8414 | 0.8514 | 0.8613 |
| *5-shot* | 0.8930 | 0.9024 | 0.9118 |
| *7-shot* | 0.9073 | 0.9131 | 0.9189 |

**Table 5:** 95 % confidence intervals for siamese Zero neural network test classification accuracy

|  | Lower limit | Mean | Upper limit |
| --- | --- | --- | --- |
| *1-shot* | 0.6266 | 0.6463 | 0.6661 |
| *3-shot* | 0.8233 | 0.8317 | 0.8402 |
| *5-shot* | 0.8762 | 0.8812 | 0.8861 |
| *7-shot* | 0.8970 | 0.9010 | 0.9050 |

**Table 6:** Tan et al. (2013) Average accuracy table (%) for different values of *k* (shots) and *d* (number of features). The highest value for each case is boldfaced.

| k | Classifier | $d = 32$ | $d = 50$ | $d = 128$ |
| --- | --- | --- | --- | --- |
| 3 | SR | **81.8** | **83.6** | N.A. |
|  | SVM | 73.5 | 74.8 | 72.4 |
|  | NS | 79.8 | 79.6 | 82.3 |
| 5 | SR | **83.9** | **87.0** | **88.6** |
|  | SVM | 75.3 | 76.8 | 77.3 |
|  | NS | 82.0 | 84.7 | 84.7 |
| 7 | SR | **85.5** | **88.2** | **89.6** |
|  | SVM | 78.7 | 80.8 | 81.6 |
|  | NS | 84.5 | 86.4 | 86.4 |

Despite Tan et al. (2013) performance results are slightly below (~ 2%) those of our best models, we consider the difference statistically and practically insignificant. Unfortunately, proper statistical hypothesis testing between our models and those of Tan et al. paper could not be carried out because confidence intervals were not reported, only average performance.

Furthermore, the variance of accuracy as a function of the number of shots (*k*) is reduced both in the bidirectional LSTM and Zero siamese network, as compared to the rest of models. This can be explained in terms of the sample complexity of few-shot learning models. In other words, the number of training samples that we need to provide (so that the learned function is within a small error range most of the time) is a function of model complexity (i.e. VC dimension) (Vapnik, 2000; Ma & Fu, 2012). Thus, relatively simple models such as fully connected networks have high bias and variance issues given they ignore the sequential aspects of birdsong phrases.

Figure 4 provides support to the *manifold hypothesis*, which states that sample complexity of the task depends only on the *intrinsic* dimension, but not the *ambient* dimension of the data manifold (Fefferman, Mitter, & Narayanan, 2016; Ma & Fu, 2012). In other words, phrase classes cluster in sub-spaces of lower dimension within the 8192-dimensional (ambient) feature space. Which means the first two principal components of the full data set may not be able to tell apart 32 phrase classes, but a larger set (*<<* 8192) could do. This may explain why the visualization in Figure 2 shows poor grouping. Maaten and Hinton argue that linear dimensionality reduction techniques such as Principal Component Analysis and classical multidimensional scaling focus on keeping the low-dimensional representations of dissimilar data points far apart. Thus, for high-dimensional data that lies on or near a low-dimensional non-linear manifold it is usually more important to keep the low-dimensional representations of very similar data points close together. (Van Der Maaten & Hinton, 2008).

In contrast, t-SNE was particularly useful at revealing cluster structure because it relied on minimizing Kullback-Leibler divergence between distributions at high and low dimension. This is unsurprising if we consider Tan et al. (2013) obtained a similar classification performance using only 128 features, which means the intrinsic dimension is low compared to that of the ambient space. Nevertheless, since t-SNE makes use of the *manifold hypothesis* by preserving local neighborhoods, in data sets with a high intrinsic dimensionality and an underlying manifold that is highly varying, the local linearity assumption on the manifold that t-SNE implicitly makes (by employing Euclidean distances between close neighbors) may be violated.

We believe the fact Zero siamese neural network, a model without non-linear transformations, performed as good as the bidirectional siamese network is best explained by the *Manifold hypothesis* in the following way: Zero network computed euclidean distances between the average of the support set per class (centroid) and each query. Then, it assigned the label of the closest centroid. Since differential features of each class were embedded in low-dimensional sub-manifolds, as shown by Figure 4 and confirmed by Tan et al. (2013) results, to classify correctly it sufficed to take the closest centroid in a non-linearly transformed 8192-dimensional feature (ambient) space.

By the other side, even if the Zero network is almost as good as the bidirectional LSTM, the bi-LSTM has a slight advantage of ~ 2% which may be greater for datasets with a different manifold structure and more classes. Moreover, since we did not carry out hyperparameter optimization for each few-shot model, it is premature to generalize any performance gain beyond *Tan2013* dataset. This is to say that our conclusions are limited to the region of the parameter space we explored, and we cannot say our models are better overall.

Finally, it is important to mention that as opposed to Tan et al. (2013), in this work we were not concerned with the effect of dimensionality reduction in linear classifiers, but only with the impact of *k* (shots) across different end-to-end siamese neural network few-shot classifiers. Overall, our results show varying degrees of sensitivity to *k* as a function of neural architectures and confirm few-shot linear models can obtain similar performance to siamese neural networks provided classes are nicely embedded in low-dimensional sub-spaces (as shown by t-SNE projection).

## 6 Main findings

We carried out a study on the capabilities of few-shot siamese neural network models for bird phrase verification and classification. From a biological perspective, our results shed light on the manifold structure and morphological distribution of *Vireo cassinii*. It known that auditory stimuli of syllables of the same class produce similar activation patterns in auditory brain areas (Koumura & Okanoya, 2016). Which means phrases that are close according to the metric learned by siamese networks, might share neural activation patterns during sensorimotor control.

Variations in the acoustic properties of birdsong are related to sound production mechanisms and features of the habitat (Derryberry, 2009). Thus, monitoring these changes across time and space can uncover causal factors of birdsong evolution and habitat selection. For instance, phrases that are close in the feature space may share a biological function or physical constraints. Further studies have to be carried out in this direction to confirm phonological similarities are meaningful at the biological level.

More broadly, this work provides a methodology for training machine learning models in class imbalance and data sparsity conditions. These are common and challenging problems in biological data (Xu & Jackson, 2019). Since they can dramatically skew the performance of classifiers by introducing a prediction bias for the majority class, addressing them is of paramount importance, specially in situations where the occurrence of false negatives is costlier than false positives (Leevy, Khoshgoftaar, Bauder, & Seliya, 2018) (Johnson & Khoshgoftaar, 2019). In our particular situation, LSTM siamese neural networks achieved state of the art performance, but we also found that computationally cheaper, and not deep-learning based models such as the *Zero Network* (which is essentially a *k*− Nearest Neighbors classifier) can achieve similar results. Nevertheless, since we did not carry out a thorough evaluation of the hyperparameter space of siamese models, it is premature to generalize any performance gain beyond *Tan2013* dataset.

By the other side, the unexpected result of *Zero network* performing as good as bi-LSTM siamese neural network raised questions about the adequacy of deep learning models. The fact that deep neural networks have been effective at more domains than simple linear classifiers does not imply complex models are always better. We think it is important to understand how deep learning models process data manifolds before drawing any conclusions in this respect. There is ongoing research leveraging methods from statistical mechanics to tackle foundational questions in this area, particularly generalization and the effects of random initialization (Bahri et al., 2020).

Furthermore, deep learning models are cognitively cheap to implement, as they are capable of learning representations without direct human input, but their computational cost is high. These computational factors, as well as those inherent to the nature of data and its distribution should be considered during the development phase of machine learning models in computational science.

Avenues not explored in this work but worth pursuing include: evaluating models beyond *Tan2013* dataset to see if learned features are universally useful across *Passeriformes* species, measuring the effect of transfer learning in classification performance, extending the model to perform segmentation and alignment of new phrases as well as classification and evaluating the stability and performance of few-shot learning models with different class groupings. Since there is little agreement as to how birdsong elements should be defined, this might be relevant from both a computational and bioacoustics point of view.

## 7 Acknowledgements

We would like to thank Dr. Caleb Rascón, Dr. Ivá n Meza and Dr. Emmanuel Martínez for reviewing this work, which was fully funded by Tecnológico de Monterrey and Consejo Nacional de Ciencia y Tecnología (CONACyT) as part of Programa Nacional de Posgrados de Calidad (PNPC) under Award Number 717358.

## 8 Appendix

**Figure 5:**
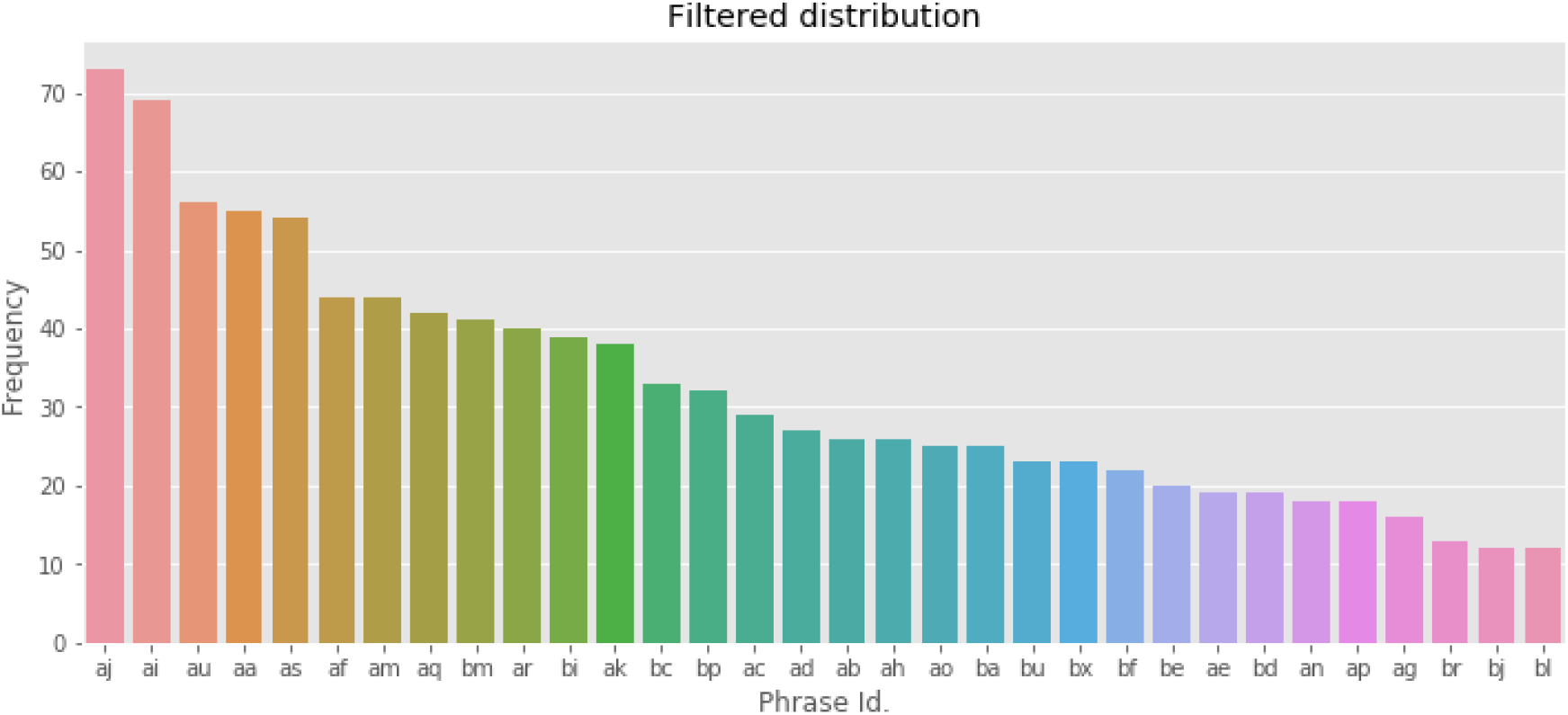
CAVI phrases with at least 12 instances

**Table 7:**
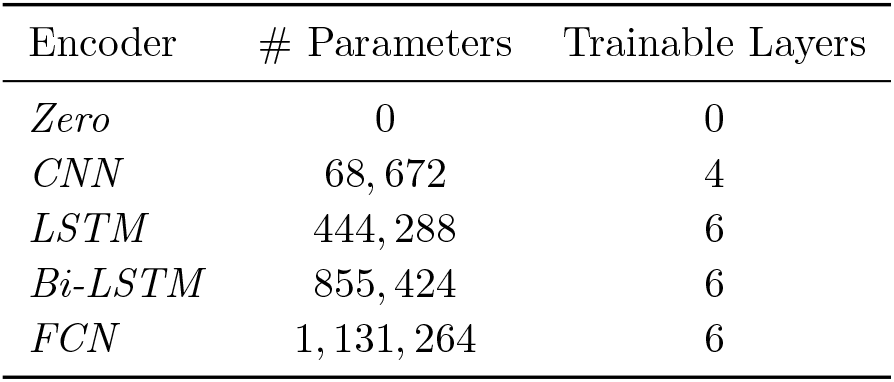
Siamese networks summary

**Figure 6:**
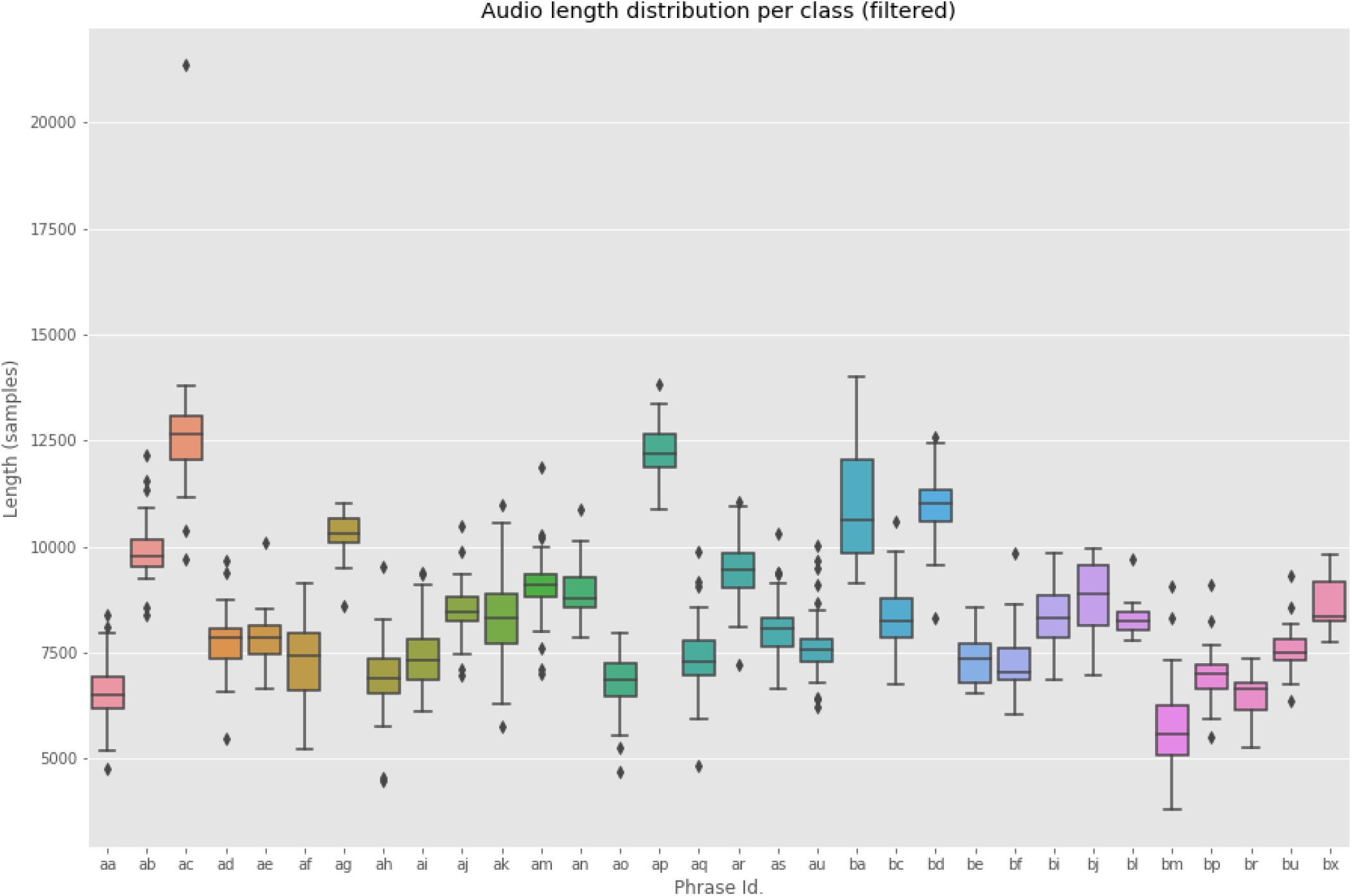
Length of CAVI phrases with at least 12 instances

**Figure 7:**
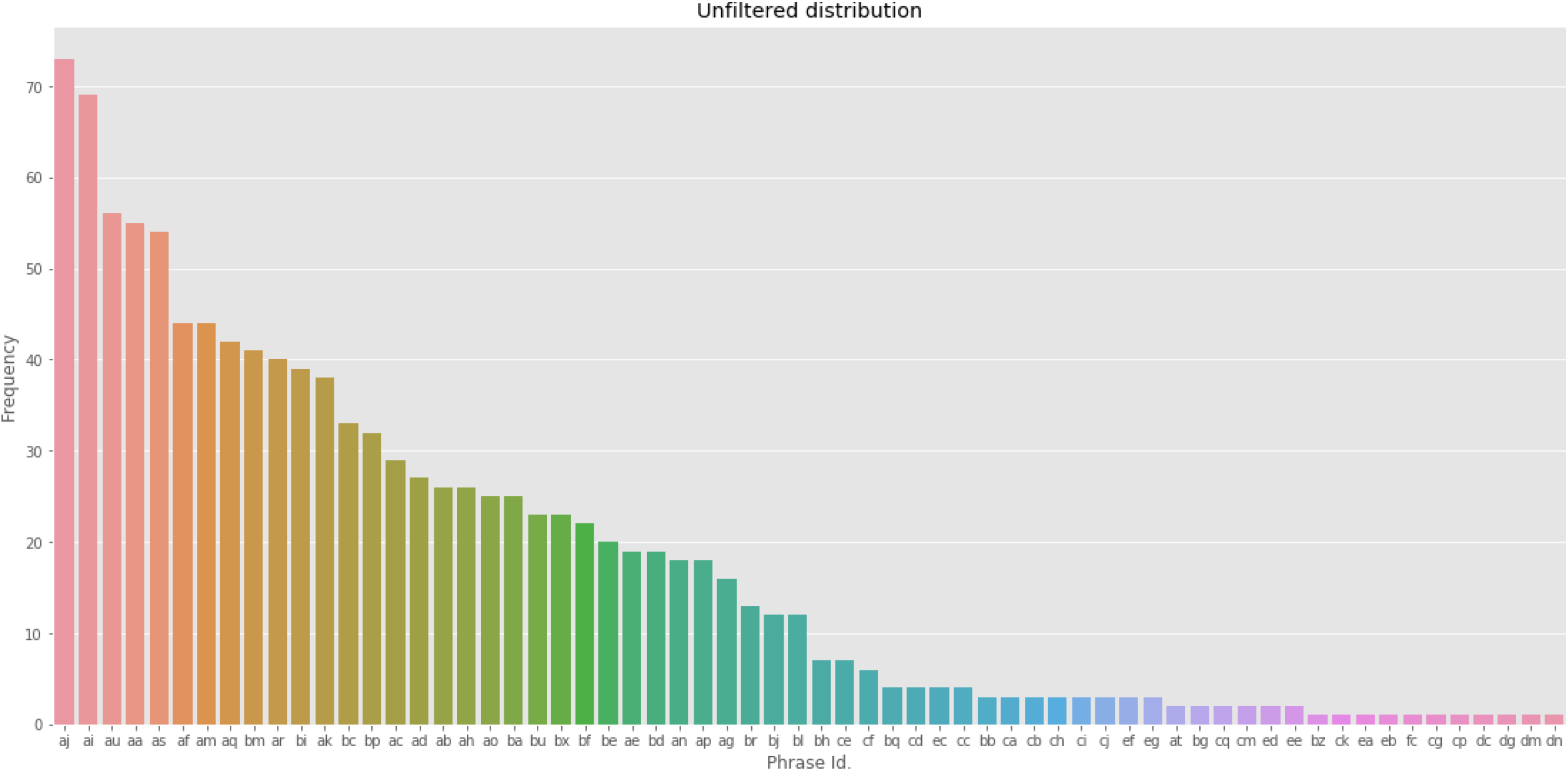
64 CAVI phrases found in *Tan2013* data set

**Figure 8:**
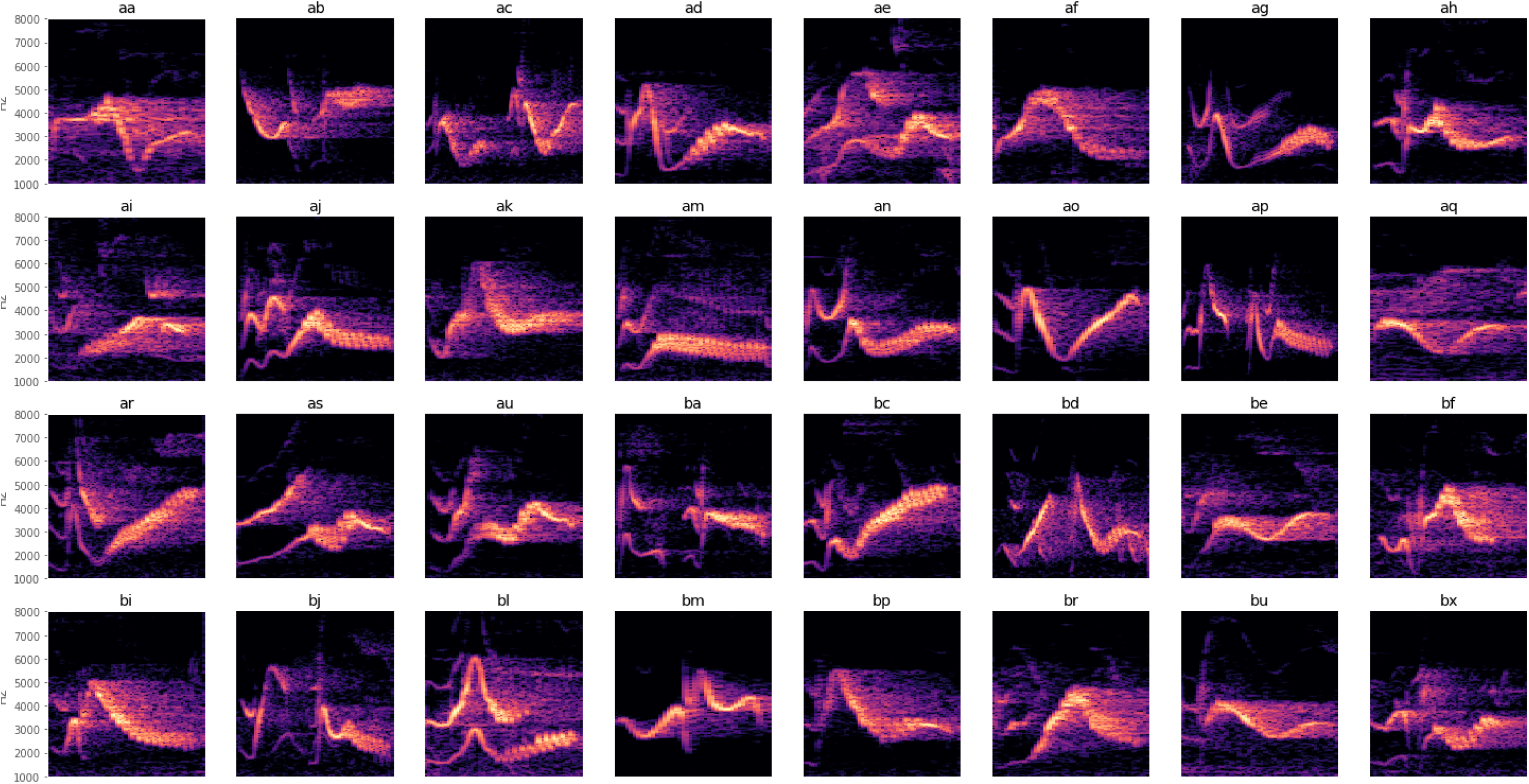
Spectrograms of CAVI phrase classes with at least 12 tokens

## References

Arriaga, J. G., Cody, M. L., Vallejo, E. E., & Taylor, C. E. (2015). Bird-db: A database for annotated bird song sequences. Ecological Informatics, 27, 21 – 25. Retrieved from http://www.sciencedirect.com/science/article/pii/S1574954115000151 doi: https://doi.org/10.1016/j.ecoinf.2015.01.007

Arriaga, J. G., Kossan, G., Cody, M. L., Vallejo, E. E., & Taylor, C. E. (2013). Acoustic sensor arrays for understanding bird communication. identifying cassin’s vireos using svms and hmms. In ECAL (pp. 827–828). MIT Press.

Bahri, Y., Kadmon, J., Pennington, J., Schoenholz, S. S., Sohl-Dickstein, J., & Ganguli, S. (2020). Statistical Mechanics of Deep Learning. Annual Review of Condensed Matter Physics. doi: 10.1146/annurev-conmatphys-031119-050745

Bioacoustics Research Program, & Program, B. R.(2014). Raven Pro: Interactive Sound Analysis Software (Version 1.5). Retrieved from http://ravensoundsoftware.com/software/raven-pro/

Boersma, P., & Weenink, D. (2011). Praat: Doing phonetics by computer. Ear and Hearing. doi: 10.1097/AUD.0b013e31821473f7

Catchpole, C. K., & Slater, P. J. B. (2008). Bird song: Biological themes and variations (Second ed.). Cambridge University Press.

Clark, C., Marler, P., & Beeman, K. (1987). Quantitative Analysis of Animal Vocal Phonology: an Application to Swamp Sparrow Song. Ethology. doi: 10.1111/j.1439-0310.1987.tb00676.x

Derryberry, E. P. (2009). Ecology shapes birdsong evolution: Variation in morphology and habitat explains variation in white-crowned sparrow song. American Naturalist. doi: 10.1086/599298

Fefferman, C., Mitter, S., & Narayanan, H. (2016, 10). Testing the manifold hypothesis. Journal of the American Mathematical Society, 29 (4), 983–1049. doi: 10.1090/jams/852

Fukuzawa, Y., Marsland, S., Pawley, M., & Gilman, A. (2017). Segmentation of harmonic syllables in noisy recordings of bird vocalisations. In International conference image and vision computing new zealand. doi: 10.1109/IVCNZ.2016.7804445

Furbee, L. (1992). Categorization and Naming in Children: Problems in Induction.:Categorization and Naming in Children: Problems in Induction. Journal of Linguistic Anthropology. doi: 10.1525/jlin.1992.2.1.120

Goguen, C. B., & Curson, D. R. (2002). Cassin’s vireo (vireo cassinii). The Birds of North America Online. Retrieved from https://doi.org/10.2173/bna.615 doi: 10.2173/bna.615

Goodfellow, I., et al. (2016). Deep learning. MIT Press.

Johnson, J. M., & Khoshgoftaar, T. M. (2019). Survey on deep learning with class imbalance. Journal of Big Data. doi: 10.1186/s40537-019-0192-5

Kaewtip, K., Taylor, C., & Alwan, A. (2016). Noise-robust hidden markov models for limited training data for within-species bird phrase classification. In Interspeech 2016 (pp. 2587–2591). Retrieved from http://dx.doi.org/10.21437/Interspeech.2016-1360 doi: 10.21437/Interspeech.2016-1360

Kershenbaum, A. (2014). Entropy rate as a measure of animal vocal complexity. Bioacoustics. doi: 10.1080/09524622.2013.850040

Kershenbaum, A., et al. (2016). Acoustic sequences in non-human animals: A tutorial review and prospectus. Biological Reviews, 91 (1), 13–52. doi: 10.1111/brv.12160

Koops, H. V., van Balen, J., & Wiering, F. (2015). Automatic Segmentation and Deep Learning of Bird Sounds. In J. Mothe et al. (Eds.), Experimental ir meets multilinguality, multimodality, and interaction (pp. 261–267). Cham: Springer International Publishing.

Koumura, T., & Okanoya, K. (2016). Automatic recognition of element classes and boundaries in the birdsong with variable sequences. PLoS ONE, 11 (7). doi: 10.1371/journal.pone.0159188

Leevy, J. L., Khoshgoftaar, T. M., Bauder, R. A., & Seliya, N. (2018). A survey on addressing high-class imbalance in big data. Journal of Big Data. doi: 10.1186/s40537-018-0151-6

Ma, Y., & Fu, Y. (2012). Manifold learning theory and applications. CRC Press.

Pearre, B., Perkins, L. N., Markowitz, J. E., & Gardner, T. J. (2017). A fast and accurate zebra finch syllable detector. PLoS ONE. doi: 10.1371/journal.pone.0181992

Planqué, B., & Vellinga, W.-P. (2005). Xenocanto. Retrieved from http://xeno-canto.org

Potamitis, I., Ntalampiras, S., Jahn, O., & Riede, K. (2014). Automatic bird sound detection in long real-field recordings: Applications and tools. Applied Acoustics. doi: 10.1016/j.apacoust.2014.01.001

Ranjard, L., & Ross, H. A. (2008). Unsupervised bird song syllable classification using evolving neural networks. The Journal of the Acoustical Society of America. doi: 10.1121/1.2903861

Tan, L. N., Alwan, A., Kossan, G., Cody, M. L., & Taylor, C. E. (2015). Dynamic time warping and sparse representation classification for birdsong phrase classification using limited training data. The Journal of the Acoustical Society of America, 137 (3), 1069–1080. doi: 10.1121/1.4906168

Tan, L. N., Kaewtip, K., Cody, M. L., Taylor, C. E., & Alwan, A. (2012). Evaluation of a sparse representation-based classifier for bird phrase classification under limited data conditions. In 13th annual conference of the international speech communication association 2012, interspeech 2012.

Tan, L. N., Kossan, G., Cody, M. L., Taylor, C. E., & Alwan, A. (2013). A sparse representation-based classifier for in-set bird phrase verification and classification with limited training data. In Icassp, ieee international conference on acoustics, speech and signal processing - proceedings. doi: 10.1109/ICASSP.2013.6637751

Van Der Maaten, L., & Hinton, G. (2008). Visualizing data using t-SNE. Journal of Machine Learning Research.

Vapnik, V. N. (2000). The Nature of Statistical Learning Theory. Springer. doi: 10.1007/978-1-4757-3264-1

Xu, C., & Jackson, S. A. (2019). Machine learning and complex biological data. doi: 10.1186/s13059-019-1689-0

Yip, K., & Sussman, G. J. (1997). Sparse representations for fast, one-shot learning. In Proceedings of the national conference on artificial intelligence (pp. 521–527). AAAI.

